# Inhibition of sodium conductance by cannabigerol contributes to a reduction of neuronal dorsal root ganglion excitability

**DOI:** 10.1101/2021.09.14.460359

**Authors:** Mohammad-Reza Ghovanloo, Mark Estacion, Peng Zhao, Sulayman Dib-Hajj, Stephen G. Waxman

## Abstract

Cannabigerol (CBG), a non-psychotropic phytocannabinoid, is a precursor for cannabis derivatives, Δ9-tetrahydrocannabinol and cannabidiol (CBD). Like CBD, CBG has been suggested as an analgesic. A previous study reported CBG (10 μM) blocks voltage-gated sodium (Nav) currents in CNS neurons. However, the manner in which CBG inhibits Nav channels, and whether this effect contributes to CBG’s potential analgesic behavior remain unknown. Genetic and functional studies have validated Nav1.7 as an opportune target for analgesic drug development. The efforts to develop therapeutic selective Nav1.7 blockers have been unsuccessful thus far, possibly due to issues in occupancy; drugs have been administered at concentrations many folds above IC_50_, resulting in loss of isoform-selectivity, and increasing off-target effects. We reasoned that an alternative approach could use compounds possessing 2 important properties: ultra-hydrophobicity and functional selectivity. Hydrophobicity could enhance absorption into neuronal cells especially with local administration. Functional selectivity could reduce likelihood of side-effects. As CBG is ultra-hydrophobic (cLogD=7.04), we sought to determine whether it also possesses functional selectivity against Nav channels that are expressed in dorsal root ganglion (DRG). We found that CBG is a ~10-fold state-dependent Nav inhibitor (K_I_-K_R_: ~2-20 μM) with an average Hill-slope of ~2. We determined that at lower concentrations, CBG predominantly blocks sodium G_max_ and slows recovery from inactivation; however, as concentration is increased, CBG also hyperpolarizes Nav inactivation curves. Our modeling and multielectrode array recordings suggest that CBG attenuates DRG excitability, which is likely linked with Nav inhibition. As most Nav1.7 channels are inactivated at DRG resting membrane potential, they are more likely to be inhibited by lower CBG concentrations, suggesting functional selectivity against Nav1.7 compared to other Navs (via G_max_ block).

## INTRODUCTION

The cannabis plant, *Cannabis sativa*, contains over 120 active constituents, which are called phytocannabinoids (1, 2). Among these phytocannabinoids, the ones that are nonpsychotropic, are of great interest for possible clinical applications. One of the key nonpsychotropic phytocannabinoids is cannabigerol (CBG) (3). CBG is a common precursor for the most widely studied cannabis derivatives, Δ9-tetrahydrocannabinol (THC) and cannabidiol (CBD) (1, 4). Although CBG has been considerably less studied than either THC or CBD, the existing literature suggests that the pharmacological properties of CBG on some receptors and targets fall in between those of THC and CBD. The most noteworthy of these targets are the human endocannabinoid receptors, CB1 and CB2. The high affinity interactions of THC at CB1 (partial agonist) and CB2 (agonist) are thought to be the underlying reason for the euphoria that is associated with THC consumption (2, 5). In contrast, CBD’s low affinity (antagonism/inverse agonism) for these receptors are suggested to cause its lack of psychoactivity (2, 5). The CBG affinity for CB receptors is reported to be between THC and CBD, and is therefore, regarded as a weak to partial agonist on these receptors (6–9).

Like CBD, several potential therapeutic applications have been suggested for CBG, which include neuroprotection against various neurological conditions such as Huntington and Parkinson diseases (10–13), as well as some gastrointestinal diseases such as colorectal cancer (14). Importantly, CBG has been implicated as an analgesic compound (15). The diversity of the range of disorders for which CBG is implicated as a potential therapeutic is also similar to the claims made for CBD (2). Unlike CBD, which has been clinically approved for use against Dravet and Lennox-Gastaut syndromes (16, 17), the use of CBG is yet to be substantiated in a human clinical trial against any excitability related conditions. CBG has been shown to modulate the activities of several ion channels, including TRPA1 (agonist), TRPV1-4 (agonist), TRIPM8 (antagonist) (9, 18, 19). Previous investigations of the cannabis-mediated blockade of voltage-gated sodium (Nav) channels have determined that CBG (10 μM) significantly decreases action potential (AP) frequency in rat CA1 hippocampal neurons and Nav current density in human neuroblastoma cells and mouse cortical neurons (20). However, the exact nature of the interactions between CBG and Nav channels, and whether these interactions could be part of CBG’s potential anti-pain/analgesic behavior remain unknown.

The transient sodium current through Nav channels initiates APs in neurons, skeletal and cardiac muscles. Any alterations to the gating properties of these channels, and subsequently the current passed through them during an AP can culminate in life-limiting conditions that could be lethal. Both increasing and decreasing of the function of Nav channels can disrupt electrical signaling (21–24).

Mutations in the primary sodium channel isoforms of the peripheral nervous system (PNS), Nav1.6-9, elicit pain and nociception-related conditions (25). Physiologically, Nav1.7 is regarded as the threshold Nav isoform that triggers APs, with Nav1.8-9 as general propagator of APs, that supersede the first spike.

Mutations in *SCN9A* that cause Nav1.7 to become hyperexcitable have been implicated in pain disorders with Mendelian inheritance patterns. This suggests that Nav1.7 has a central role in pain response in humans. Dominant gain-of-function (GOF) mutations in Nav1.7 are found in patients with paroxysmal extreme pain disorder (PEPD) (26) and inherited erythromelalgia (IEM) (27). Conversely, recessive mutations in Nav1.7 have been linked with complete insensitivity to pain (CIP) (28).

CIP, in part, has resulted in Nav1.7 being regarded as a highly validated target for analgesic drug development, as CIP mutations impart little to no effects on motor, cardiac, or cognitive functions (28–30). However, much of the extensive efforts that have gone into the development of selective blockers of Nav1.7 have been unsuccessful in vivo, possibly due to challenges in drug occupancy of Nav1.7. Getting around the problems with receptor occupancy, has relied on drug administration many folds above the IC_50_, which result in loss of selectivity, and hence, off-target effects (29).

From a theoretical perspective, some of the aforementioned issues may be remedied if a compound possesses 2 important properties. First, the compound should be ultra-hydrophobic, and second, it should display functional selectivity. The compound hydrophobicity could be crucial, in that it could enhance absorption into the fatty neuronal cells, upon a properly carried out local administration. The functional selectivity could reduce likelihood of unwanted side-effects. CBG with a calculated LogD of 7.04 fits the hydrophobicity criteria. In this study, we sought to determine whether CBG also fits the second criteria, with respect to functionally targeting of dorsal root ganglion (DRG) excitability, at least in part, via Nav inhibition. As there are some similarities in the physicochemical properties of CBG and CBD, and because CBD has been thoroughly described with respect to Nav interactions (2, 31–34), we used the prior CBD studies to guide our investigations of CBG activity in the present paper.

## RESULTS

### CBG is a state-dependent inhibitor of sodium currents

Our first goal in this study was to determine the concentration-response of CBG on Nav currents, and to find out whether there is state-dependence. To do this, we performed voltage-clamp of stably transfected Nav1.7 in HEK cells. We used a protocol to examine state-dependent inhibition across a range of holding-potentials where channel inactivation varied (35). We first held channels at a holding-potential of −110 mV where channels are almost all in the restingstate, while pulsing the channels 180 times at 1 Hz to allow CBG to reach equilibrium. Then, we depolarized the holding-potential by 10 mV two more times and repeated the pulse train at each voltage (**Fig. 1A**). To construct the concentration-response, individual cells were exposed to single concentrations of CBG, then normalized inhibition at each concentration was pooled and fit with a Hill Langmuir equation, providing IC_50_ (4.6-18.8 μM) and Hill-slopes. We show the fractional block of sodium currents from the last pulse from each holding-potential (**Fig. 1B**). Representative current traces are shown in **Fig. 1A**. Interestingly, the sodium current inhibition has very steep Hill-slopes of 1.2-2.3 across all three holding potentials (**Fig. 1B**). This indicates that CBG, similar to CBD (32), likely does not inhibit Navs through a 1:1 binding mechanism. This suggests that multiple interactions contribute to CBG inhibition.

**Figure 1.**
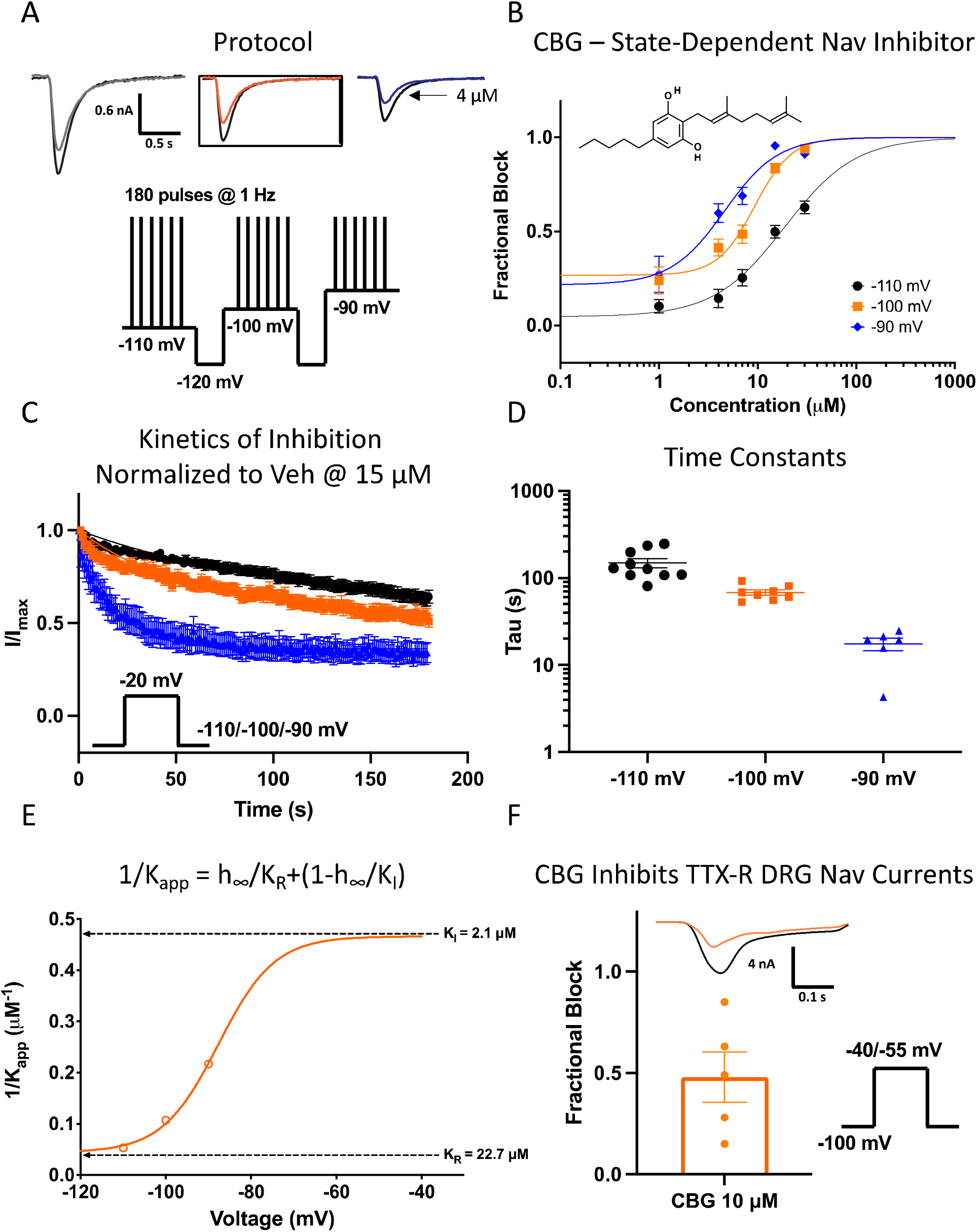
State-dependence of CBG as a voltage-dependent sodium channel inhibitor. (**A**) Pulse protocol showing 180 pulses run at 1 Hz at each holding-potential and representative current traces. (**B**) CBG potency at three holding-potentials at pulse 180 (3 min) in Nav1.7 (IC_50_ (μM): −110 mV = 18.8 ± 0.6, −100 mV = 9.3 ± 0.7, −90 mV = 4.6 ± 0.7; n = 9-19). Structure of CBG is shown at the top left. (**C**) Kinetics of inhibition of Nav1.7 at the noted holding potentials (Mean (s): −110 mV = 148.3 ± 18.2, −100 mV = 68 ± 4.6, −90 mV = 17.5 ± 2.9; n = 9-19). (**D**) distribution of kinetics time constants. (**E**) Apparent Kd at different voltages was well fit with a 4-state model invoking different potencies for resting and inactivated-state block. (**F**) Manual voltage-clamp of primary rat DRG neurons (mean fractional inhibition = 0.48 ± 0.1; n = 5).

Next, we investigated CBG’s kinetics of Nav1.7 inhibition by measuring peak INa amplitude over 3 minutes during a series of pulses to −20 mV from the same three holding potentials mentioned above. The observed rates of equilibration (time constant observed, ⍰_obs_) of inhibition were measured by fitting single exponential decays to inhibition of currents. The fraction of inhibition was normalized against the response in vehicle and plotted against the time elapsed after CBG (15 μM) addition, which was set at t = 0 (**Fig. 1C-D**). We found that the rate at which 15 μM CBG inhibits Nav1.7 becomes faster as the holding potential gets depolarized. This potential-dependent change in inhibition kinetics is congruent with the IC_50_ relationships in **Fig. 1B** and is consistent with the idea of CBG being a state-dependent Nav inhibitor.

**Fig. 1E** shows a plot of the inverse of the apparent IC_50_ fit with a 4-state binding model that used parameters obtained from the Boltzmann fit of the voltage-dependence of steady-state inactivation (SSI) from a 500 ms pre-pulse (33, 36, 37). The potency numbers were based on the results shown in **Fig. 1B**. This established that the apparent potency is directly related to the proportion of inactivated channels at different holding-potentials. These results demonstrate that CBG inhibits the sodium current from both rest (~23 μM) and inactivated-states (~2 μM); however, the potency of CBG is about ~11-fold greater for inhibiting inactivated compared to resting-states (**Fig. 1E**). The magnitude fold difference in apparent potency of CBG between the inactivated- to resting-states of Nav is comparable to CBD, which displays a state-dependence that is closer to ~10-fold (32).

It is previously shown that CBD has no molecular selectivity among the channels in the Nav superfamily (31–33). The equipotent inhibition of the Navs has been in part ascribed to the binding of CBD at the central cavity-fenestration interface within the lipid phase of the membrane. Because this site of the Nav structure is a conserved part of the pore-domain, then interactions at this site would be structurally non-selective in nature (2, 31). To determine, whether CBG shares this lack of structural selectivity, we voltage-clamped isolated rat DRG neurons (held at −100 mV and pulsed to −40 or −55 to elicit maximal current). The bath solutions contained 300 nM tetrodotoxin (TTX), to block out all the TTX-sensitive Nav channels (IC_50_ of TTX-sensitive channels is ~10-30 nM (33, 38)). Then, we perfused 10 μM CBG for ~3 minutes (time-matched to the experiments shown in Nav1.7-HEK cells in **Fig. 1B**). Then, we measured peak amplitude differences in the TTX-resistant Nav current in response to CBG (**Fig. 1F**). Our results suggested that 10 μM CBG, on average, inhibited ~50% of the TTX-resistant current in rat DRG, which is in close agreement with the data in **Fig. 1B**. This suggests that like CBD, CBG likely lacks structural selectivity among Navs.

### CBG prevents Nav opening, but does not affect activation voltage-dependence

We next examined the effects of CBG on Nav activation by measuring peak channel conductance at membrane potentials between −120 and +30 mV. We show the effects of 15 μM CBG on peak conductance as a function of membrane potential (**Fig. 2A**). About ~90% of the sodium conductance was inhibited at 15 μM CBG. Next, we plotted the maximal sodium conductance (G_max_) from 1-30 μM CBG at −25 mV (potential at which maximal amount was elicited). Our results indicate that CBG imparts a concentration-dependent decrease in G_max_ (**Fig. 2B**). This decrease started to become statistically significantly different from vehicle at 4 μM (p<0.05).

**Figure 2.**
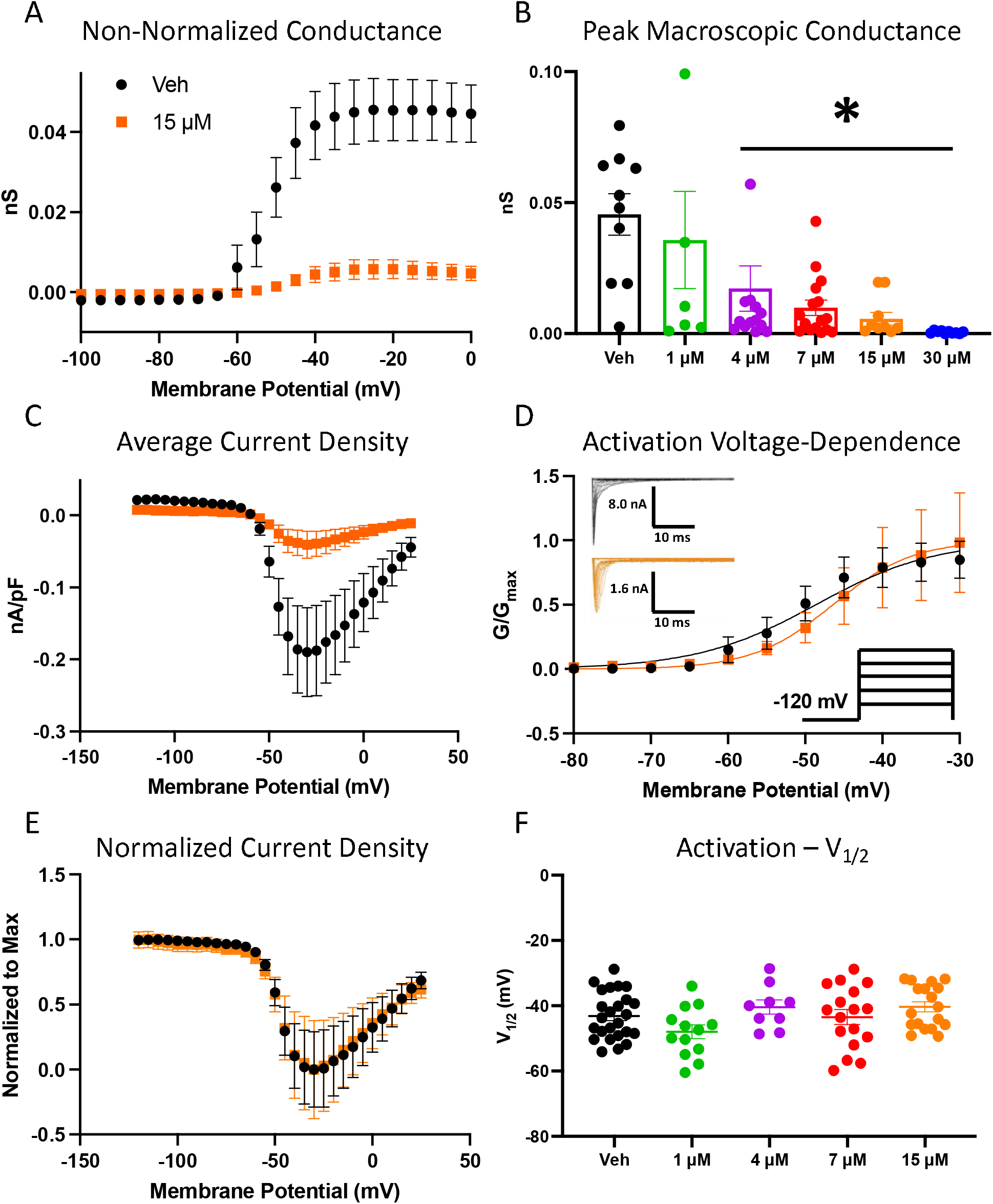
CBG Does not alter activation and inhibits conductance in Nav1.7. (**A**) Conductance difference in Nav1.7 in vehicle and 15 μM CBG as a function of membrane potential. (**B**) Quantification of peak macroscopic conductance at −25 mV across different CBG concentrations (Veh = 0.046 ± 0.008; 1 μM = 0.036 ± 0.02; 4 μM = 0.017 ± 0.009; 7 μM = 0.0099 ± 0.003; 15 μM = 0.0057 ± 0.002; 30 μM = 0.0002 ± 0.0002; n = 8-17). (**C**) Average current density of hNav1.7 in vehicle and 15 μM CBG. (**D**) Voltage-dependence of activation as normalized conductance plotted against membrane potential (Vehicle: V_1/2_ = −48.6 ± 0.6 mV, Slope = 7.7 ± 0.6, n = 10; CBG: V_1/2_ = −46.4 ± 0.3 mV, Slope = 5.1 ± 0.3, n = 10). (**E**) Normalized current density, further displaying unaltered activation. (**F**) Midpoints of activation across concentrations (in mV) (Veh = −43 ± 1; 1 μM = −48 ± 2; 4 μM = −40 ± 2; 7 μM = −44 ± 2; 15 μM = −40 ± 2; n = 8-17).

In **Fig. 2C** we show a plot of sodium current density as peak INa divided by membrane capacitance (nA/pF) as a function of membrane potential, which consistent with the G_max_, show a ~90% reduction in magnitude at 15 μM. The normalized conductance is plotted against membrane potential (**Fig. 2D**), showing that neither 15 μM nor any of the other CBG concentrations induce large changes in the midpoint (V_1/2_) or apparent valence (slope, k) of activation of the available Nav channels (p>0.05) (**Fig. 2D-F, Fig. S1A**). Therefore, exposures to CBG concentration-dependently prevents Navs from conducting; however, these exposures do not alter the voltage-dependence of activation. This effect is similar to what has been reported for CBD (32).

### CBG concentration-dependently hyperpolarizes the inactivation curve, but does not alter open-state inactivation

We next measured the voltage-dependence of inactivation from a 500 ms pre-pulse duration. Generally, durations that are in the range of low hundreds of milliseconds are considered more implicative of fast inactivation than slow inactivation. However, 500 ms may be considered to trigger an intermediate amount of inactivation. Indeed, measuring inactivation using longer pre-pulses would be more physiologically relevant in Nav1.7 which predominantly resides in cells where the resting membrane potential (RMP) is by in large depolarized compared to the channel availability V_1/2_. The normalized current amplitudes at the test-pulse are shown as a function of pre-pulse voltages (**Fig. 3A-C**). Our results suggested that while there is a concentration-dependent shift in the inactivation curves, the shifts started to become statistically significant from vehicle at 15 μM (**Fig. 3D**). We also found that the slopes of the inactivation curves at none of the concentrations were significantly different from vehicle (p>0.05) (**Fig. S1B**). The current at the test-pulse was inhibited by ~90% at 15 μM; however, unlike activation, the voltage-dependence of SSI of the remaining current was hyperpolarized by ~12 mV (p<0.0001). This indicates CBG increased the propensity for channels to inactivate over the 500 ms pre-pulse in channels that were not inhibited from opening from rest, further suggesting that CBG stabilizes the inactivated states of Nav channels. CBG’s overall effect of hyperpolarizing the inactivation curves is similar to what is reported for CBD.

**Figure 3.**
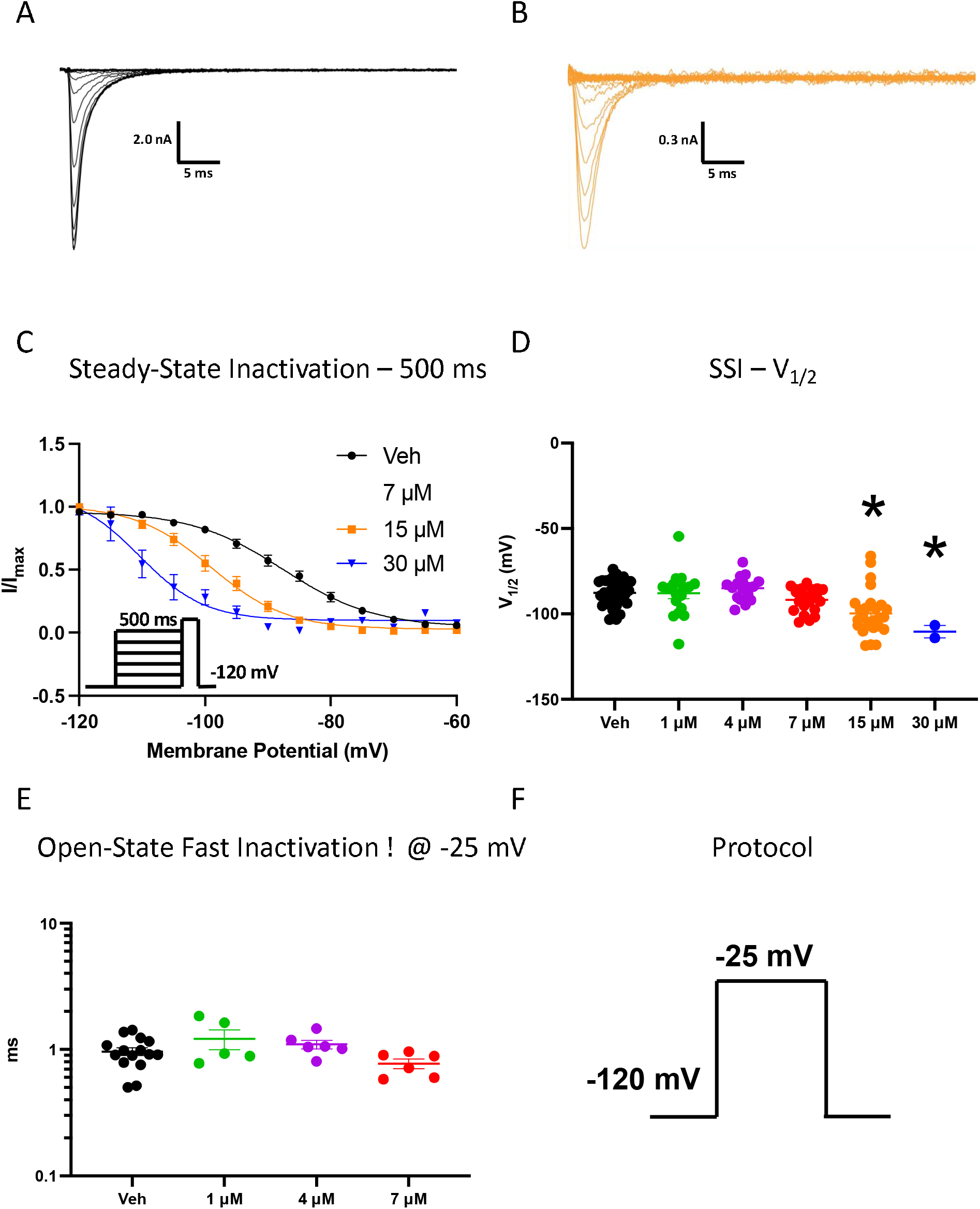
CBG hyperpolarizes inactivation curve in Nav1.7. (**A**-**B**) Sample macroscopic current traces of veh and 15 μM CBG. (**C**) Voltage-dependence of 500 ms inactivation as normalized current plotted against membrane potential fit with Boltzmann. (**D**). (**E**) Quantification of SSI midpoints (in mV) (Veh = −88 ± 1; 1 μM = −88 ± 1; 4 μM = −85 ± 3; 7 μM = −92 ± 2; 15 μM = −100 ± 3; 30 μM = −110 ± 4; n = 2-33). (**E**) Mean open-state fast inactivation time constants (ms) (Veh = 0.96 ± 0.07; 1 μM = 1.2 ± 0.2; 4 μM = 1.1 ± 0.1; 7 μM = 0.8 ± 0.1; n = 5-15). (**F**) The protocol that was used for (**E**).

We also measured the rate time constant of open-state inactivation at −25 mV (also known as true fast inactivation) using trace fitting by an exponential function, which did not differ significantly at any of the CBG concentrations compared to vehicle (p>0.05) (**Fig. 3E-F**). CBG’s lack of interaction with the open-state of the channel is also similar to CBD (32), and is consistent with the proposed blocking scheme for an ultra-hydrophobic compound, which suggests that as the drug becomes more hydrophobic, it tends to interact more with resting and inactivated states of the channel (2).

### CBG stabilizes inactivated-states of Nav channels by slowing recovery

To assess the time-dependence and degree of stabilization of the inactivated-state we then measured the recovery from inactivation of Nav1.7 in different CBG concentrations (1-30 μM). This was done after depolarizing pre-pulse durations of 20 ms (fast inactivated), 500 ms (intermediate inactivated), or 5 s (slow inactivated), from a holding-potential of −120 mV. To measure recovery from inactivation, we held the channels at −120 mV to ensure that the channels were fully available, then pulsed the channels to −20 mV at one of the above durations and allowed different time intervals at −120 mV to measure recovery as a function of time (**Fig. 4A**). The mean normalized recovery following the pre-pulse in vehicle and various CBG concentrations were plotted and fit with a bi-exponential function (**Fig. 4B-D**).

**Figure 4.**
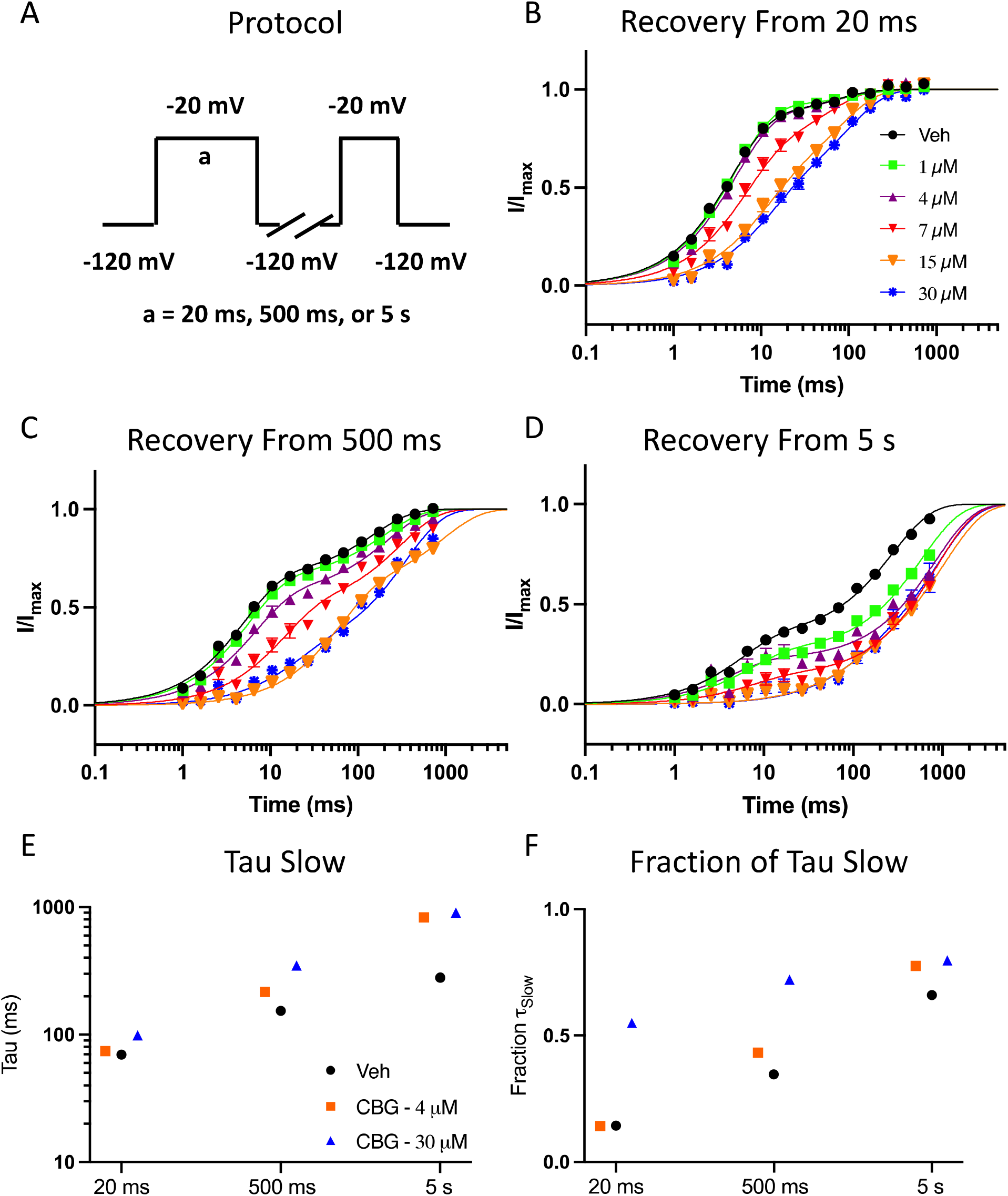
CBG Slows recovery from inactivation in Nav1.7. (**A**) Shows the protocol that was used to measure CBG effect on channel recovery from duration pre-pulses. (**B**) Recovery from inactivation in the presence of 0 (Veh)-30 μM CBG, from 20 ms. (**C**) 500 ms. (**D**) 5s. (**E-F**) The slow components of recovery from inactivation in vehicle and CBG at 20 ms, 500 ms, and 5 s are shown on the left Y-axis, and the fraction of slow to fast component of recovery from inactivation is shown on the right Y-axis.

Our results suggested that when the channels are fast inactivated, the lower concentrations of 1-4 μM do not alter the time-course of recovery compared to vehicle, and only at 7 μM can a slowing of recovery become detectable. However, at the high CBG concentrations of 15-30 μM, even 20 milliseconds of inactivation accumulation is sufficient to significantly slow channel recovery (**Fig. 4B**). When the pulse duration is increased to 500 ms, we see that both of lower concentrations, notably 4 μM, begin to display a slowing of the recovery (**Fig. 4C**). Expectedly, after 5 seconds of inactivation accumulation, even the lowest concentration of 1 μM causes a major slowing of recovery (**Fig. 4D**). This is an important finding, as the inactivation V_1/2_ Nav1.7 is hyperpolarized relative to the RMP of DRG neurons, and therefore, at RMP (when there is no AP firing), much of the membrane-bound Nav1.7 channels are in inactivated state, which would be more reminiscent of the data shown in **Fig. 4D** than **Fig. 4B**, which suggests that under physiological conditions, relatively lower concentrations of CBG would prevent Nav1.7 channels from opening. To further illustrate this point, we show the ⍰_Slow_ and fraction of the recovery fit with ⍰_Slow_ plotted in **Fig. 4E-F**, which show that CBG increases the fraction of recovery that is slow and the time constant of the slow component of recovery from inactivation from all three pre-pulse durations. This further indicates that CBG slows the recovery from inactivation supporting the idea that CBG stabilizes the inactivated-states of the channel; however, it does so at lower concentrations (e.g., 4 vs. 30 μM) than would be expected based on steady-state measurements shown in **Fig. 3C-D**. This could have major physiological implications.

### CBG inhibits conductance more potently than it hyperpolarizes inactivation

The fundamental differences between reducing conductance vs. left-shifting availability have important mechanistic pharmacological consequences. A compound that reduces G_max_, reduces the number of available channels from opening; conversely, a compound that left-shifts inactivation, increases the likelihood of the channels that do open, to inactivate. Although both mechanisms contribute to a loss of excitability, a reduced G_max_ may be preferable to prevent AP firing in the first place. As shown in **Fig. 2-3**, the significant effects of CBG’s on G_max_ and inactivation V_1/2_, started to occur at different concentrations. To quantify this difference between the two effects, we first subtracted mean number across CBG concentrations from the mean number from vehicle from **Fig. 2B** (G_max_) and **Fig. 3D** (Inactivation V_1/2_). This calculation provided the Δ between CBG effects at a given concentration versus vehicle. Then, we divided the Δ for each of G_max_ and V_1/2_ by the vehicle numbers to obtain the Δpercentage of vehicle. The results from these calculations are plotted in **Fig. 5A**, which show that CBG alters the Nav G_max_ at lower concentrations compared to the inactivation V_1/2_. Interestingly, by the time 15 μM CBG is reached, only ~10% of the vehicle G_max_ is left.

**Figure 5.**
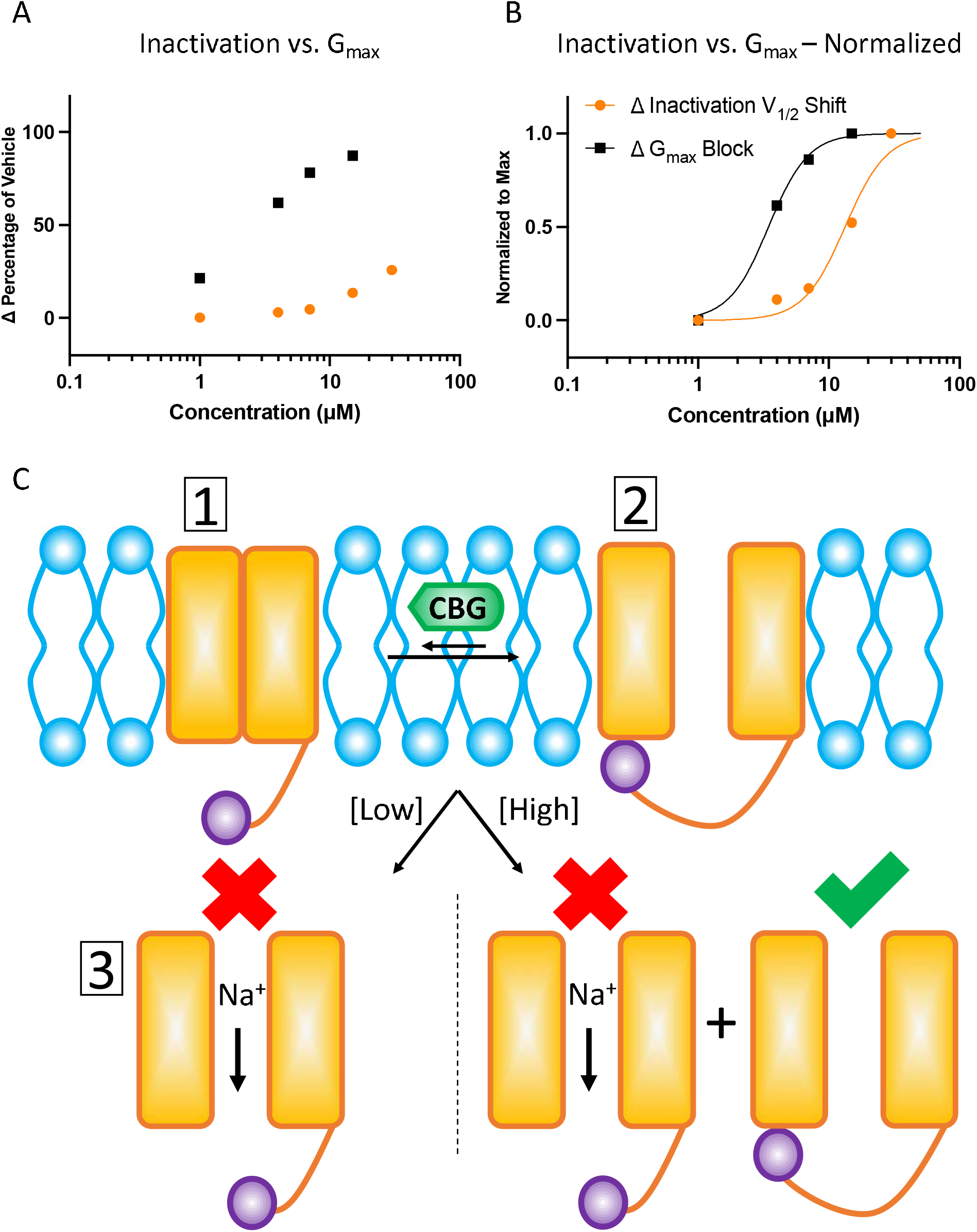
CBG Effects on conductance versus inactivation. (**A**) Comparison of concentration-dependent effects of CBD on G_max_ vs. inactivation as a percentage of vehicle. (**B**) Normalized relationship of the info from (**A**) and fit with the Hill equation (Inactivation = 13.3 ± 1.0, G_max_ =3.4 ± 1.0). (**C**) Cartoon representation of the concentration-dependent modality of CBG effects on Nav channels.

Next, to obtain an approximate potency difference between the two effects, we normalized the numbers in **Fig. 5A**, to the maximal values, which are plotted in **Fig. 5B**. This normalization was based on a logical assumption of physiological pharmacology. Although Δpercentage shift of V_1/2_ at the maximal concentration of 30 μM is only at about 25% of vehicle, because 15-30 μM almost completely abolish G_max_, then any further shifts in V_1/2_ would be physiologically inconsequential. In other words, if no sodium current is left, an increased likelihood of inactivation would be irrelevant. When we fitted the data with the Hill equation, we found that CBG’s IC_50_ in inhibiting G_max_ is approximated to 3.5 μM, with the IC_50_ for V_1/2_ being estimated to 13.2 μM. As higher concentrations of CBG are likely to culminate in modulation of diverse targets, we suggest that CBG’s potential therapeutic value in reducing Nav activity with respect to pain would likely stem from inhibiting G_max_, and not hyperpolarization of the inactivation curves.

Overall, our results on the empirical effects of CBG on Nav channels suggest that due to its high hydrophobicity (a higher partitioning/distribution of CBG in lipid vs. water indicates a greater preference for the membrane phase at equilibrium), CBG readily gets inside the membrane, from where it interacts with the resting- and inactivated-states of Nav channels, though with a greater affinity for the channels that are in the inactivated conformations. If the RMP is such that a given Nav channel is chronically accumulating inactivation (e.g., Nav1.7 in DRG), then lower levels of CBG in the membrane would be sufficient to keep channels from opening, which would culminate in cellular reduction of excitability (**Fig. 5C**).

### CBG reduces neuronal excitability in a Hodgkin-Huxley model of DRG

To test the effects of CBG on neuronal excitability, we used a modified version of the Hodgkin-Huxley model to simulate a DRG neuron’s excitability (39, 40). We ran two simulations of the CBG condition, at 2 and 15 μM. As the RMP in the DRG is estimated to ~-60 mV, based on the 4-state model in **Fig. 1E**, we determined that 2 μM CBG would prevent opening of ~30% of Nav channels. Therefore, in the 2 μM simulation, we only reduced the overall sodium conductance, by about 30%. In the 15 μM simulation, we reduced the sodium conductance by 87% and hyperpolarized the inactivation curve by 12 mV, which is based on **Fig. 3C-D**. In our simulations, the channels were given a series of step-wise current injections with increasing intensities at each step for 100 ms. Each 100 ms step was followed by a 50 ms recovery period in which no current injection was applied (**Fig. 6A-B**). Our results suggest that the peak amplitude of the first AP in 2 μM CBG is smaller than vehicle. This is consistent with the reduction of peak sodium conductance caused by CBG. Consistent with the massive effects of 15 μM on conductance and inactivation, the resulting simulations displayed big reductions of excitability across stimulus injection intensities.

**Figure 6.**
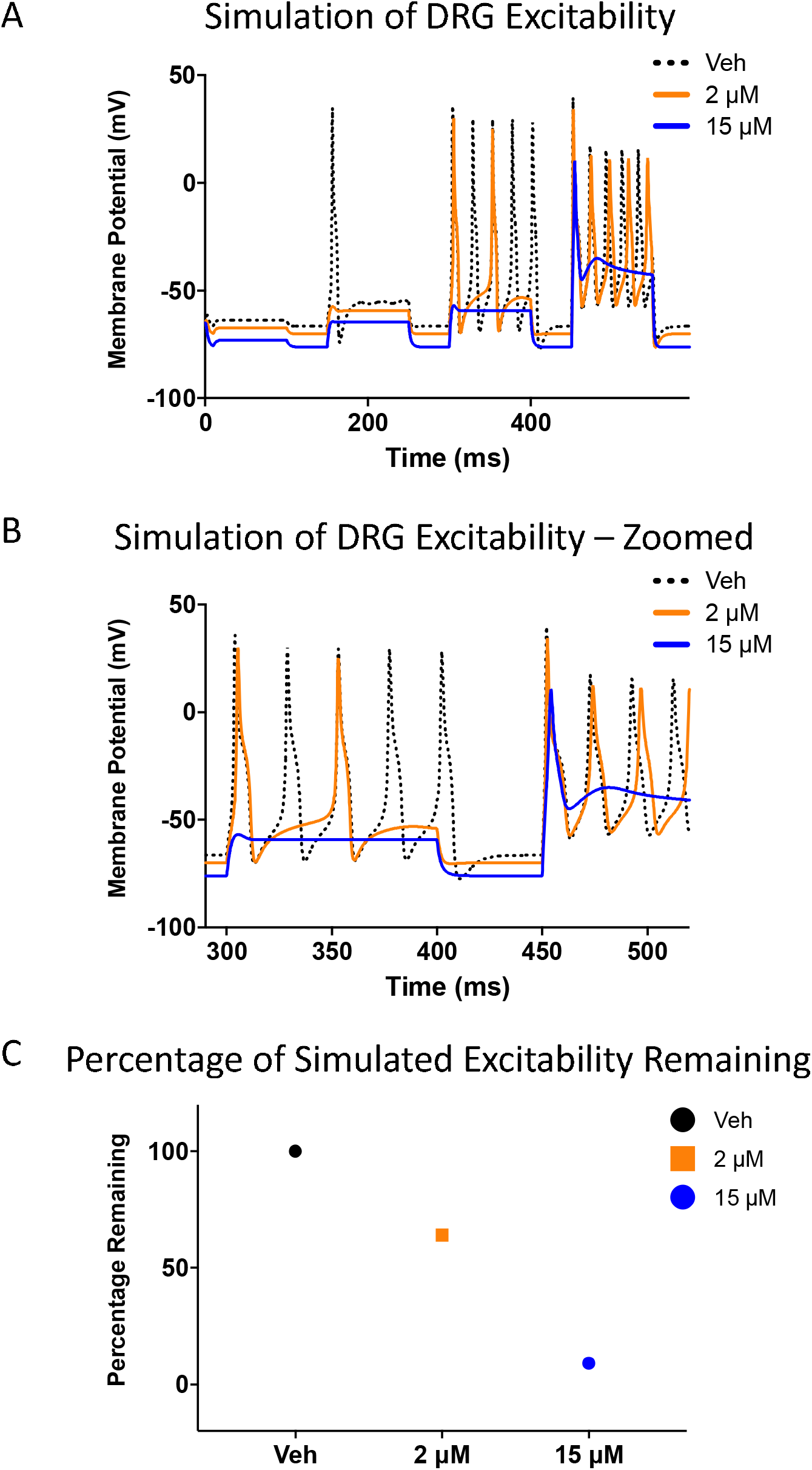
CBG Reduces excitability in an action potential model. (**A**) Simulation of the effects of CBG on action potential morphology over a series of increasing current injection intensities. (**B**) Zoomed-in simulation of action potentials from the first interval shown in (**A**). (**C**) Number of peaks in CBG divided by number of peaks in vehicle.

The overall effect of CBG on the AP morphology is that at all current injection intensities, CBG reduces the number of APs, leading to a net loss of excitability. To further quantify these simulation predictions, we normalized the number of spikes in CBG conditions to vehicle, which suggested that 2 μM reduces APs by 36%, and 15 μM reduces APs by 91% (**Fig. 6C**).

### CBG reduces spontaneous steady-state excitability in rat DRG neurons

To assess whether the predicted CBG-mediated reduction in neuronal excitability holds, we measured excitability of rat DRG neurons using multielectrode array recordings (MEA). We measured spontaneous firing over a 10-minute period in MEA wells that had vehicle, 2, and 15 μM CBG (**Fig. 7A-B**) (the activity of all wells was compared before vehicle/compound addition, and there was no significant difference among the wells). Our results suggested that CBG concentration-dependently reduced neuronal firing. Normalization of the mean reduced excitability of CBG conditions to the vehicle suggested that 2 and 15 μM CBG reduced firing by 32% and 89%, respectively (**Fig. 7C**). While these numbers are in close agreement with the simulation results, CBG most likely hits other targets besides Navs, particularly at 15 μM, which collectively result in reduced DRG firing. These findings suggest that CBG at low micromolar concentrations, at least in part via Nav channels, can reduce DRG firing, which can potentially be used against pain.

**Figure 7.**
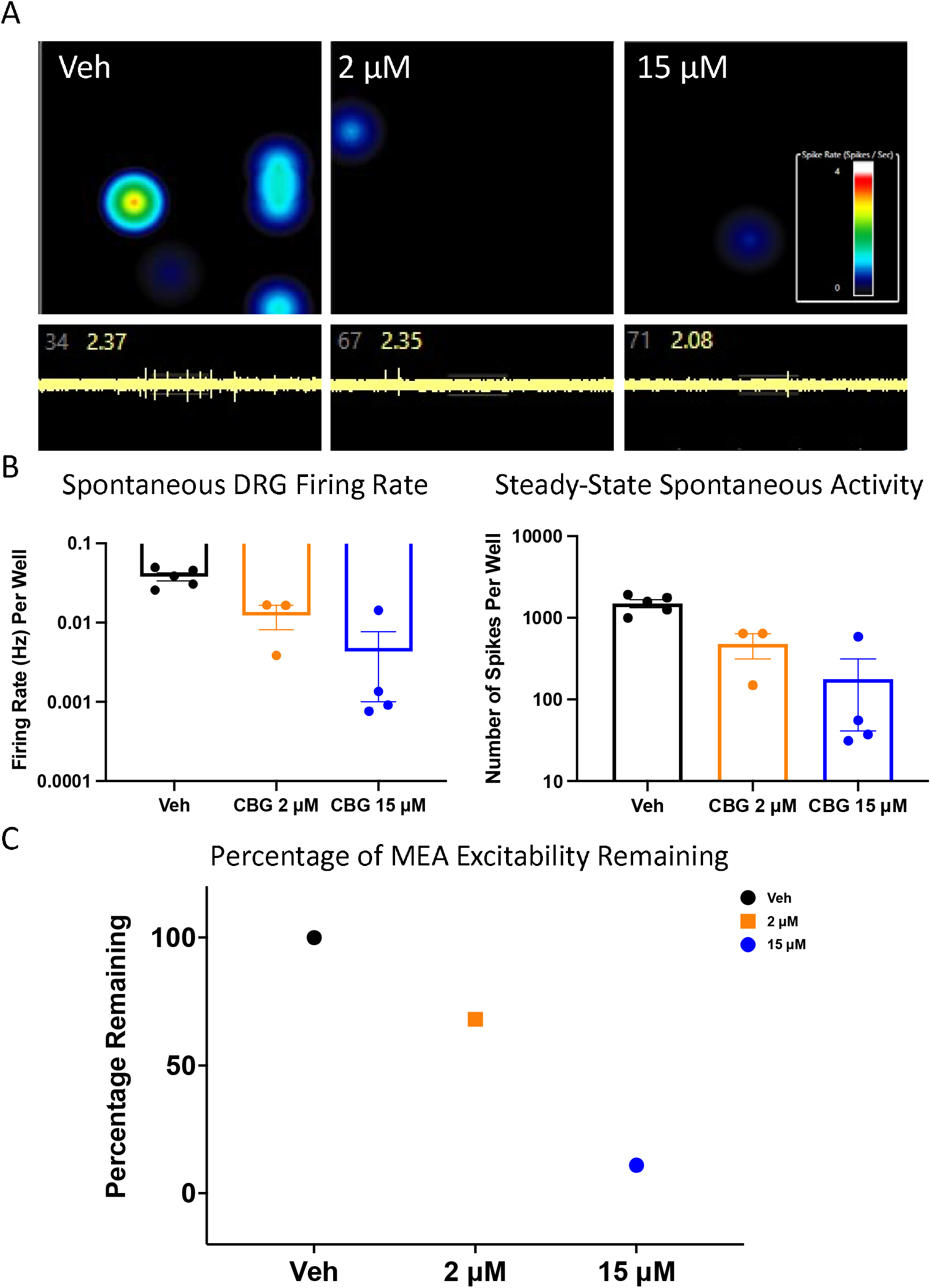
CBG Reduces excitability of primary rat DRG neurons in MEA. (**A**) Representative images of MEA recordings of AP firing at vehicle, 2 and 15 μM CBG. The firing frequency of each active electrode is color coded: white/red mean high, and blue/black mean low frequencies. (**B-C**) Quantification of MEA data showing firing rate and steady-state spontaneous activity of the primary neurons. (**A**). (**C**) The DRG excitability in CBG is normalized to vehicle.

## DISCUSSION

Cannabis derivatives have a long history of being used as therapeutics (1, 2, 4). The recent clinical success of CBD against seizure disorders proves that some of the historical anecdotal claims about cannabinoids have real therapeutic merits. Although in recent years CBD has been extensively studied from many angles, the related compound, CBG, is still far from being thoroughly understood. Despite having a higher affinity for the CB receptors than CBD, CBG’s interactions at these sites are not sufficiently strong to impart psychoactivity. This in itself, merits further investigations into this compound. Although CBD’s interactions with Nav channels as a mechanism of clinical efficacy against convulsions remains speculative, the crucial role of Nav channels in excitability, and the potency at which CBD inhibits Navs suggests an important relationship between the two effects (2). Of course, the CBD and Nav relationship, though likely vital, is not the only important relationship. This is because many studies have also shown CBD modulates many other targets including other ion channels, among other receptors (32, 41–44). Furthermore, the CBD relationship with Navs provides a critical blueprint to gain potential understanding of how CBG could produce the acclaimed therapeutic effects, notably analgesia (15).

### CBG vs. CBD from a Nav perspective, and why CBG may be a better option for pain

In this study, we set out to determine whether CBG could be used as a potential viable compound to functionally target the excitability of DRG neurons. Because of the reports of CBG being an analgesic agent, and as it is a very hydrophobic compound, we argued that CBG could feasibly reach the DRG neurons, upon local administration (45). As Nav channels are the ignitors of excitability in DRGs (and other excitable cells) (38), we first characterized CBG’s effects on Nav1.7. Our findings suggested that CBG’s general manifestations on Navs are very similar to CBD; however, there are some key differences. For instance, although both CBD and CBG display ~10-fold state-dependence, CBG inhibits Nav channels about 2-fold less potently than CBD, from both resting and inactivated-states. This indicates that CBG has a higher ratio of concentration per fold state-dependence than CBD (**Table 1**). This suggests that there is a net increased amount of CBG needed to impart a net loss of excitability than CBD, which for a hydrophobic compound could implicate about a 2-fold reduction in possible toxicity. This means that low Δmicromolar changes in free CBG concentration (e.g., ~1-5 μM), produces smaller effects on other targets than CBD does. For instance, if we assume ~2 μM CBG (**Fig. 1E, Fig. 7C**) is sufficient to reduce DRG firing initiation, then ~20 μM higher than the desired concentration of 2 μM would be needed to reduce ~50% of Nav1.4 in the skeletal muscle. Whereas for CBD, only ~10 μM higher than the desired concentration of 1 μM (32) would produce similar toxicity on Nav1.4 in muscle. It is worth mentioning that at such high concentrations of either compound, much more pronounced cytotoxicity is likely to happen on several other targets, before other Navs would be hit (2). Additionally, as CBG is more hydrophobic than CBD, it can get absorbed into bio-membrane more readily, and possibly have a longer lasting effect than CBD.

**Table 1.** The comparison of CBG to CBD with respect to Navs. Parameters with asterisk were taken from literature (32).

| **Parameter** | **CBG** | **CBD** |
| --- | --- | --- |
| LogD | 7.04 | 6.32 |
| State-dependence | ~10 | ~10* |
| IC<sub>50</sub> on Nav channels (μM) | ~2-22 | ~1/12* |
| Concentration / fold state-dependence (Ratio) | 2 | 1.1 |

Our voltage-clamp data in rat DRG neurons suggested that, like CBD, CBG most likely lacks structural selectivity among Navs. However, given that CBG is a state-dependent inhibitor (along with its high hydrophobicity), we suggest that DRGs could be among its high affinity cellular targets. This is because despite having only 9 members (excluding Nav2.1), the Nav superfamily orchestrate diverse and complex patterns of AP generation. The activity of Navs is determined by the local membrane potential they are subjected to. Each one of the 9 members of the family has a unique, though sometimes similar to one another, availability V_1/2_. If the V_1/2_ of each Nav is compared with the RMP of the cell it predominantly resides in, a rough estimate of Nav inactivation at RMP could be determined. In **Fig. 8**, we show a comparison of 5 general tissue-specific cell types with RMPs and V_1/2_ of their respective predominantly-expressed Navs, which are taken from the literature (33, 46–55). These comparisons suggest that in the 5 cell types (other cells and tissue must be taken into consideration on an individual basis) that are shown, Nav1.7 relative to its native DRG environment is the most inactivated of the 9 subtypes (**Fig. 8A-E**). This would suggest that any state-dependent Nav inhibitor that lacks structural subtype-selectivity, would have a higher functional selectivity for Nav1.7 (at low micromolar concentrations), in the DRG. This could in theory suggest a reduction in the probability of toxicity in other tissues (e.g., in the heart). With respect to CBG, this general principle could work in concert with its high hydrophobicity to reduce pain initiation (**Fig. 8F**). This could explain the reduction of spontaneous firing of DRG neurons in our MEA data. However, if CBG is to be considered as a viable anti-pain therapeutic, mode of administration, bioavailability, and tissue distribution must be considered.

**Figure 8.**
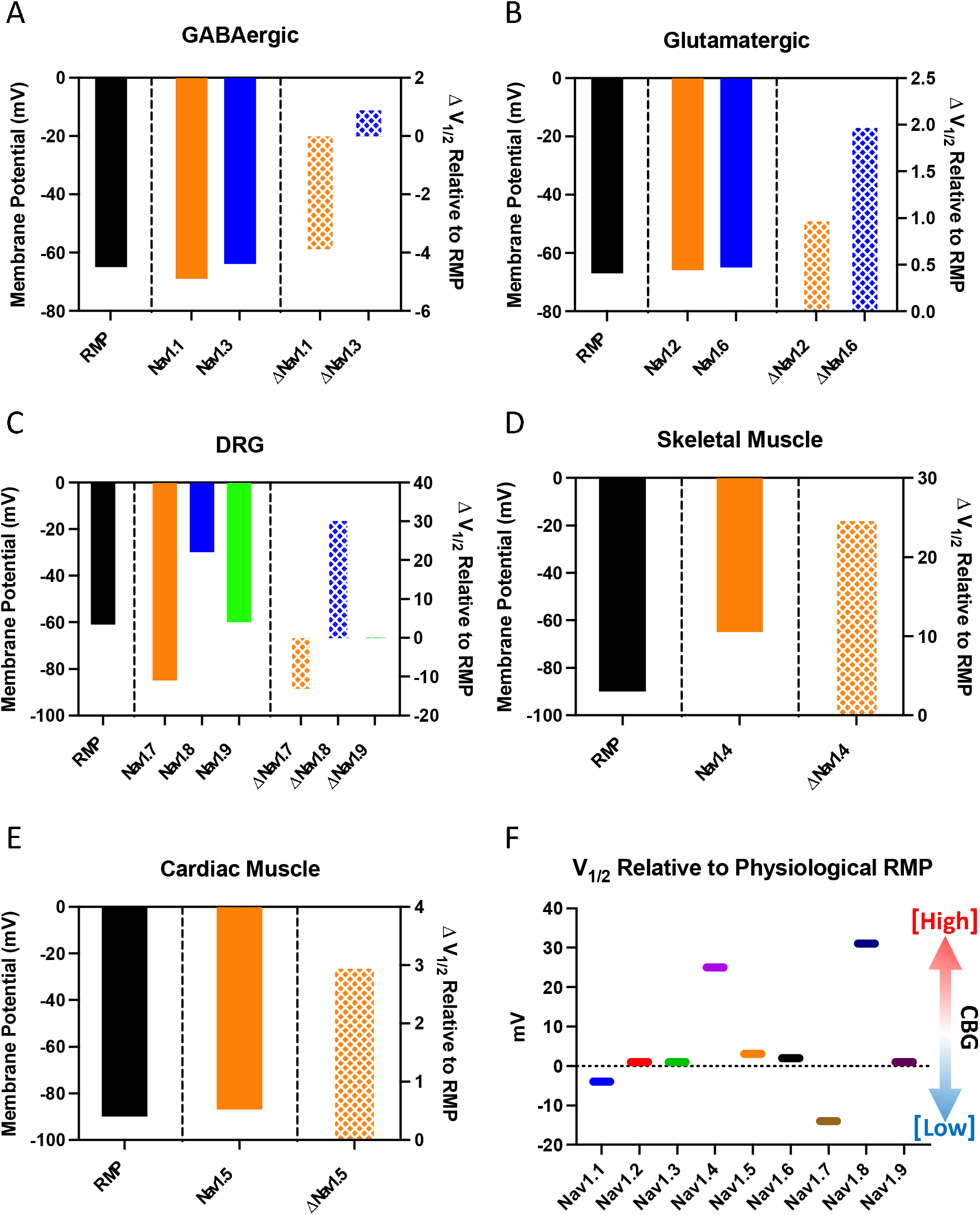
Comparison of Nav channel availabilities in some native tissues, where they are predominantly found. (**A**) GABAergic. (**B**) Glutamatergic. (**C**) DRG. (**D**) Skeletal muscle. (**E**) Cardiac muscle. (**F**) Nav availability relative to the resting membrane potential of the tissues described in this figure.

### CBG – possible mechanism of Nav inhibition

A summary of our results with a proposed mechanism of action is shown in the cartoon in **Fig. 5C**. The similarities in the general effects of CBG and CBD on Nav gating and kinetics prompt us to suggest that CBG could bind at a similar location within the Nav fenestration-central cavity site that CBD binds (31). Additionally, the steepness of the concentration-response curves along with the physicochemical properties of CBG could also suggest a possible modulation of the membrane elasticity, which could contribute to the hyperpolarization of the inactivation curve at higher than physiologically conceivable concentrations (e.g., >15 μM), without altering voltage-dependence of activation. This effect would be similar to what is suggested for amphiphilic compounds (56, 57), and CBD (33), which are thought to modulate the biophysical properties of the membrane, at concentrations that are many times over what is required to modulate high-affinity targets. Indeed, CBG’s potency across the described targets in the literature suggest that its high affinity targets are modulated within sub-micromolar to low micromolar ranges (9, 18, 19). This is another reason for why we suggest CBG’s anti-pain via Navs would also be in the low micromolar range.

In conclusion, our findings in this paper characterize the effects of CBG on Navs and suggest that these interactions could be crucial to the compound mediated analgesia.

## MATERIALS & METHODS

### Cell culture

Human Embryonic Kidney (HEK-293) cells were used for automated patch-clamp experiments. HEK-293 cells were stably transfected. The human *SCN1B* cDNA construct was co-transfected into each cell line. All cells were incubated at 37°C/5% CO_2_.

Primary Sensory Neuron Isolation for MEA. Animal studies followed a protocol approved by the Yale University and Department of Veterans Affairs West Haven Hospital Animal Use Committees.

Each well of 12-well MEA plate (Axion Biosystems) was coated with poly-D-lysine (50 μg/ml) and laminin (10 μg/ml). DRGs from rat pups (P0-P5) were harvested, and neurons were dissociated as described previously (58). Briefly, DRGs were incubated for 20-min at 37°C in complete saline solution (CSS) [in mM: 137 NaCl, 5.3 KCl, 1 MgCl2, 25 sorbitol, 3 CaCl2, and 10 N-2-hydroxyethylpiperazine-N′-2-ethanesulfonic acid (Hepes), adjusted to pH 7.2 with NaOH] containing in 1.5 mg/ml Collagenase A (Roche) and 0.6 mM EDTA, followed by a 20-min incubation at 37°C in CSS containing 1.5 mg/ml Collagenase D (Roche), 0.6 mM EDTA, and 30 U/mL papain; DRGs were then triturated in 0.5 mL of DRG media [DMEM/F12 with 100 U/ml penicillin, 0.1 mg/ml streptomycin (Invitrogen), 2 mM L-glutamine, and 10% fetal bovine serum (Hyclone)] containing 1.5 mg/mL BSA (low endotoxin) and 1.5 mg/mL trypsin inhibitor (Sigma). After trituration, cell suspension was filtered with 70-μm cell strainer (Becton Dickinson). The cell suspension was centrifuged (100 g for 3 min), and the cell pellet was transfected with 1.0 μg of cheriff eGFP using a Nucleofector IIS (Lonza, Basel, Switzerland) and Amaxa Basic Neuron SCN Nucleofector Kit (VSPI—1003) with SCN Basic Neuro Program 6. Transfected neurons were diluted with DRG media and seeded 60 μl per well. After neurons were settled for 50 min, each well was fed with 1.44 ml of DRG media (final volume 1.5 ml) supplemented with nerve growth factor (50 ng/mL) and glial cell line-derived neurotrophic factor (50 ng/mL) and maintained at 37°C in a 95% air/5% CO2 (vol/vol) incubator for three days before MEA recording.

Primary Sensory Neuron Isolation for patch clamp. Same enzymatic procedure as for DRG neuron isolation for MEA, except that after trituration, 100 μl of cell suspension was seeded directly onto each poly-D-lysine/laminin-coated coverslips (BD), and incubated at 37°C in a 95% air/5% CO2 (vol/vol) incubator. After 45 min for neurons to attach to the coverslips, DRG media was added into each well to a final volume of 1.0□ml and the neurons were maintained at 37°C in a 95% air/5% CO2 (vol/vol) incubator for xx hours before patch-clamp recording.

### Automated patch-clamp

Automated patch-clamp recording was used for all HEK experiments. HEK cell lines stably expressing the full-length cDNAs coding Nav1.7 α-subunit were generated. The human β1-subunit was co-expressed in our internally-generated cell lines. GenBank accession number for α-subunits was Nav1.7 (NM_002977). Sodium currents were measured in the whole-cell configuration using a Qube-384 *(Sophion A/S, Copenhagen, Denmark)* automated voltage-clamp system. Intracellular solution contained (in mM): 120 CsF, 10 NaCl, 2 MgCl_2_, 10 HEPES, adjusted to pH7.2 with CsOH. The extracellular recording solution contained (in mM): 145 NaCl, 3 KCl, 1 MgCl_2_, 1.5 CaCl_2_, 10 HEPES, adjusted to pH7.4 with NaOH. Liquid junction potentials calculated to be ~7 mV were not adjusted for. Currents were low pass filtered at 5 kHz and recorded at 25 kHz sampling frequency. Series resistance compensation was applied at 100% and leak subtraction enabled. The Qube-384 temperature controller was used to manipulate recording chamber temperature for all experiments, set to 22 ± 2 °C at the recording chamber. Appropriate filters for cell membrane resistance (typically >500 MOhm), Series resistance (<10 MOhm) and Nav current magnitude (>500 pA at a test pulse from a resting HP of −120 mV) were routinely applied to exclude poor quality cells. Vehicle controls were run on each plate to enable correction for any compound independent decrease of currents over time. Baselines were established after 20 minutes in vehicle. Fractional inhibition was measured as current amplitude from baseline to maximal inhibition after 20-minute exposure to test compound unless otherwise noted. Normalized mean inhibition data were fit to the Hill-Langmuir equation:

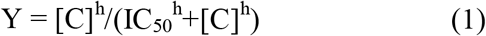

to estimate the half maximal inhibitory concentration (IC_50_ value); where Y is the normalized inhibition, C the compound concentration, IC_50_ the concentration of test compound to inhibit the currents 50%, and h the Hill coefficient. Data analysis was performed using Analyzer (Sophion A/S, Copenhagen, Denmark) and Prism (GraphPad Software Inc., La Jolla, CA, USA) software.

### Manual patch-clamp

Whole-cell patch-clamp recordings were performed using same solution as above. All recordings were made using an EPC-10 patch-clamp amplifier (HEKA Elektronik, Lambrecht, Germany) digitized at 20 kHz via an ITC-16 interface (Instrutech, Great Neck, NY, USA). Voltage-clamping and data acquisition were controlled using PatchMaster/FitMaster software (HEKA Elektronik, Lambrecht, Germany) running on a Windows computer. Current was low-pass-filtered at 10 kHz. Leak subtraction was performed automatically by software using a P/4 procedure following the test pulse. Giga-ohm seals were allowed to stabilize in the on-cell configuration for 1 min prior to establishing the whole-cell configuration. Series resistance was less than 5 MΩ for all recordings. Series resistance compensation up to 80% was used when necessary. Before each protocol, the membrane potential was hyperpolarized to −100 mV to ensure complete availability of TTX-R currents. All experiments were conducted at 22 ± 2 °C. Analysis and graphing were done using FitMaster software (HEKA Elektronik) and Igor Pro (Wavemetrics, Lake Oswego, OR, USA).

### Compound preparation

CBG was purchased from Cayman Chemicals. Powdered CBG was dissolved in 100% DMSO to create stock. The stock was used to prepare drug solutions in extracellular solutions at various concentrations with no more than 0.5% total DMSO content. TTX was prepared in water, and also diluted in extracellular solutions.

### Activation protocols

To determine the voltage-dependence of activation, we measured the peak current amplitude at test pulse potentials ranging from −120 mV to +30 mV in increments of +5 mV for 500 ms. Channel conductance (G) was calculated from peak I_Na_:

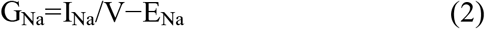

where G_Na_ is conductance, I_Na_ is peak sodium current in response to the command potential V, and E_Na_ is the Nernst equilibrium potential. Calculated values for conductance were fit with the Boltzmann equation:

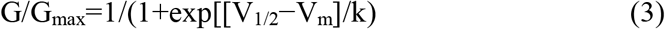

where G/G_max_ is normalized conductance amplitude, V_m_ is the command potential, V_1/2_ is the midpoint voltage and k is the slope.

### Steady-state inactivation protocols

The voltage-dependence of fast-inactivation was measured by preconditioning the channels from −120 to +30 mV in increments of 5 mV for 500 ms, followed by a 10 ms test pulse during which the voltage was stepped to −20 mV. Normalized current amplitudes from the test pulse were fit as a function of voltage using the Boltzmann equation:

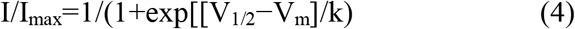

where I_max_ is the maximum test pulse current amplitude.

### State-dependence protocols

To determine state-dependence, potency was measured from three different holding-potentials (−110, −100, −90 mV). The protocol started with a holding-potential of −110 mV followed by 180 × 20 ms depolarizing pulses to 0 mV at 1 Hz. Then, the holding-potential was depolarized by 10 mV, and the 180-pulse protocol was repeated until −90 mV was reached.

### Recovery from inactivation protocols

Recovery from inactivation was measured by holding the channels at −120 mV, followed by a depolarizing pulse to 0 mV, then the potential was returned to −120 mV. This was followed by a depolarizing 10 ms test pulse to 0 mV to measure availability. Recovery from inactivation was measured after pre-pulse durations of 20 ms, 500 ms, and 5 s and fit with a bi-exponential function of the form:

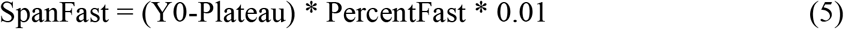

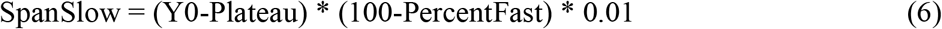

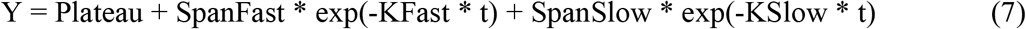

Where t is time in seconds, Y0 is the Y intercept at t=0, KFast and KSlow are rate constants in units the reciprocal of t, PercentFast the fraction of the Y signal attributed to the fast-decaying component of the fit.

### Kinetics of inhibition

The kinetics of CBG block were measured at three potentials at 15 μM. The channels were held at respective holding potentials followed by pulses to −20 mV. The blocked sodium current was normalized to vehicle and subsequently fit with a single exponential function:

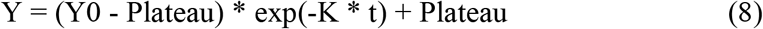

### Action potential modeling

Neuronal action potential modeling was based on a modified Hodgkin-Huxley model (39, 40). The model was modified to match the properties of DRG cells (39, 40). The modified parameters were based on electrophysiological results obtained from whole-cell patch-clamp experiments in this study. The model accounted for activation voltage-dependence, steady-state inactivation voltage-dependence, and peak sodium currents.

### Multielectrode array recordings

Multielectrode array (MEA) experiments were performed with a multi-well MEA system (Maestro, Axion Biosystems) according to a recently developed protocol (59). DRGs were dissociated and cultured on MEA plates, maintained at 37°C in a 5% CO_2_ incubator. A 12-well recording plate was used, embedded with a total of 768 electrodes. For each experiment, multiple wells were used to assess rat DRGs.

### Statistics

A t-test or ANOVA (if appropriate) were used to compare the mean responses. All statistical p-values report the results obtained from tests that compared experimental conditions to the control conditions. A level of significance α=0.05 was used with p-values less than 0.05 being considered to be statistically significant. All values are reported as means ± standard error of means (SEM) for n recordings/samples.

---

**Figure S1.**
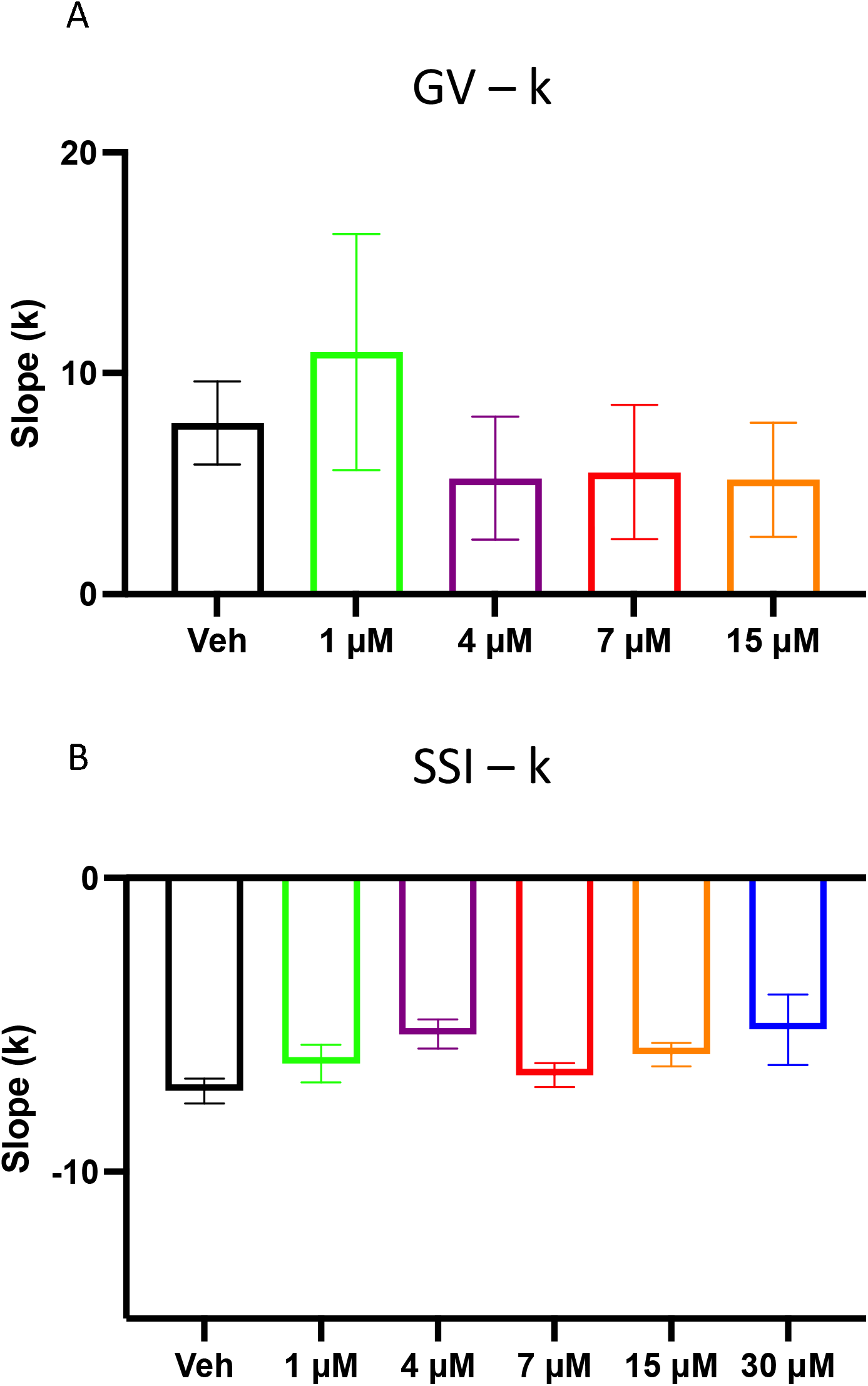
Slopes of activation and inactivation curves. (**A**) GV. (**B**) SSI.

